# Variant Graph Craft (VGC): A Comprehensive Tool for Analyzing Genetic Variation and Identifying Disease-Causing Variants

**DOI:** 10.1101/2023.12.12.571335

**Authors:** Jennifer Li, Andy Yang, Benedito A Carneiro, Ece Gamsiz Uzun, Lauren Massingham, Alper Uzun

## Abstract

The Variant Call Format (VCF) file is providing a structured and comprehensive text file that contains essential information about variant positions in the genome, as well as other critical details, such as alleles, genotype calls, and quality scores. Due to its rich data and format, the VCF file has become an increasingly popular resource for researchers and clinicians alike, enabling them to interpret and understand genomic variation data. However, analyzing and visualizing these files poses significant challenges, demanding access to diverse resources and a robust set of features for in-depth exploration. We introduce Variant Graph Craft (VGC), a VCF file visualization and analysis tool offering a wide range of features for exploring genetic variations, including extraction of variant data, intuitive visualization of variants, and the provision of a graphical representation of samples, complete with genotype information. Furthermore, VGC seamlessly integrates with external resources to offer valuable insights into gene function and variant frequencies in sample data. VGC offers gene function and pathway information from Molecular Signatures Database (MSigDB) for GO terms, as well as KEGG, Biocarta, Pathway Interaction Database, and Reactome. Additionally, the tool also provides a dynamic link to gnomAD for variant information, and includes ClinVar data for pathogenic variant information. VGC operates locally, assuring users of data security and privacy by eliminating the need for cloud-based VCF uploads. It supports the Human Genome Assembly Hg37, ensuring compatibility with a wide range of data sets. With its versatility, VGC accommodates various approaches exploring genetic variation data, and can be tailored to the specific needs of the user by using optional phenotype input data. In conclusion, VGC is a useful resource for exploring genetic variation in a secure and user-friendly environment. With its user-tailored set of features, this tool enables researchers and clinicians to easily explore and understand genomic variation data in a comprehensive and accessible manner. From identifying specific genetic mutations to analyzing patterns of variation across the genome, the VCF file visualization and analysis tool is an essential tool for those working in the field of genomics. VGC is freely available at https://sites.brown.edu/gmilab/variantgraphcraft/

## Introduction

In recent years, advancements in genome sequencing technologies have enabled researchers to generate vast amounts of genomics data. However, with this flood of information comes the need for tools that can analyze and visualize this data effectively. One of the key challenges in analyzing genetic data is dealing with the complexity and the size of variant data stored in VCF (Variant Call Format) files. These files contain information about genetic variations, including single nucleotide polymorphisms (SNPs), insertions, deletions, and structural variations. Analyzing and visualizing VCF files can be challenging task, as it requires accessing several different resources and using multiple features to explore variations. The conventional way of visualizing VCF files involves using command-line tools, which can be challenging for non-experts to use. While the existing variant call format (VCF) file visualizing tools offer useful summaries and some interactive functionalities, there are still several challenges and limitations associated with them. One of the major challenges is scalability, as many of the existing tools may struggle to handle large-scale datasets efficiently. This can lead to potential issues with performance and data processing, especially when dealing with large and complex datasets. The development of free software tools such as vcflib, bio-vcf, cyvcf2, hts-nim, and slivar offers a range of solutions for processing the VCF (Variant Call Format), potentially addressing scalability issues in handling large datasets [1]. Another limitation of these tools is the lack of or limited interactivity, as many of them do not provide dynamic and interactive environments for exploring variant data. This can make it difficult for researchers to fully understand and analyze the data and explore potential associations between genetic variants and phenotypes. In addition, some of the existing VCF file visualizing tools can be confusing to use and may require significant expertise to operate effectively. Some tools have too many dependencies based on the origin of the programming language and new updates may crash the program, which can add to the complexity of using these tools. Furthermore, compatibility issues may arise due to the different VCF file formats used by different tools, which can make it difficult to compare results between different tools. To address these challenges and limitations, several user-friendly VCF file visualization and analysis tools have been developed that offer a wide range of features for visualizing genetic variations and exporting filtered data. In the field of genomic research, there are several well-known bioinformatics tools that significantly enhance data analysis and visualization capabilities. These include IGV (Integrative Genomics Viewer), which offers an interactive platform for genomic datasets visualization [2]; VCF-Server, tailored for managing and querying Variant Call Format (VCF) files [3]; VCF.Filter, allowing for the intricate filtering of VCF files [4]; and BrowseVCF, providing a user-friendly interface for VCF file exploration [5]. Additionally, GEMINI (Genome Exploration and Mining INteractive Interface) focuses on the integrative analysis and variant prioritization within VCF files [6]. VCFtools is a comprehensive package for manipulating and interpreting VCF files, including data comparison, summarization, and statistical analysis [7]. VIVA is designed for the intuitive visualization and analysis of genomic variants, facilitating complex data interpretation through a graphical interface [8]. Together, these tools form a robust suite for genomic data management, analysis, and visualization, catering to a variety of research needs in the genomics field. However, despite the improvements made, there is still room for further enhancements to improve scalability, customizability, interactivity, complexity, and compatibility. To overcome these limitations, we have developed VGC, a VCF analysis and visualization tool designed to extract and visualize variant data from VCF files with multiple customizable options. VGC offers a comprehensive range of features for visualizing and analyzing the genomic data, empowering users to conduct their analysis locally on their machines or servers. VGC is specifically designed to efficiently handle VCF files and provide users with a flexible and user-friendly environment for conducting variant analysis. With VGC, researchers can now explore the genetic variations and gain insights into gene function and variant frequency in different populations, providing a valuable resource for advancing genetic research.

## Materials and Methods

VGC is a tool designed for analyzing variant data and visualizing VCF files. It utilizes a range of technologies and libraries to offer a user-friendly experience (Figure 1).

**Figure 1.**
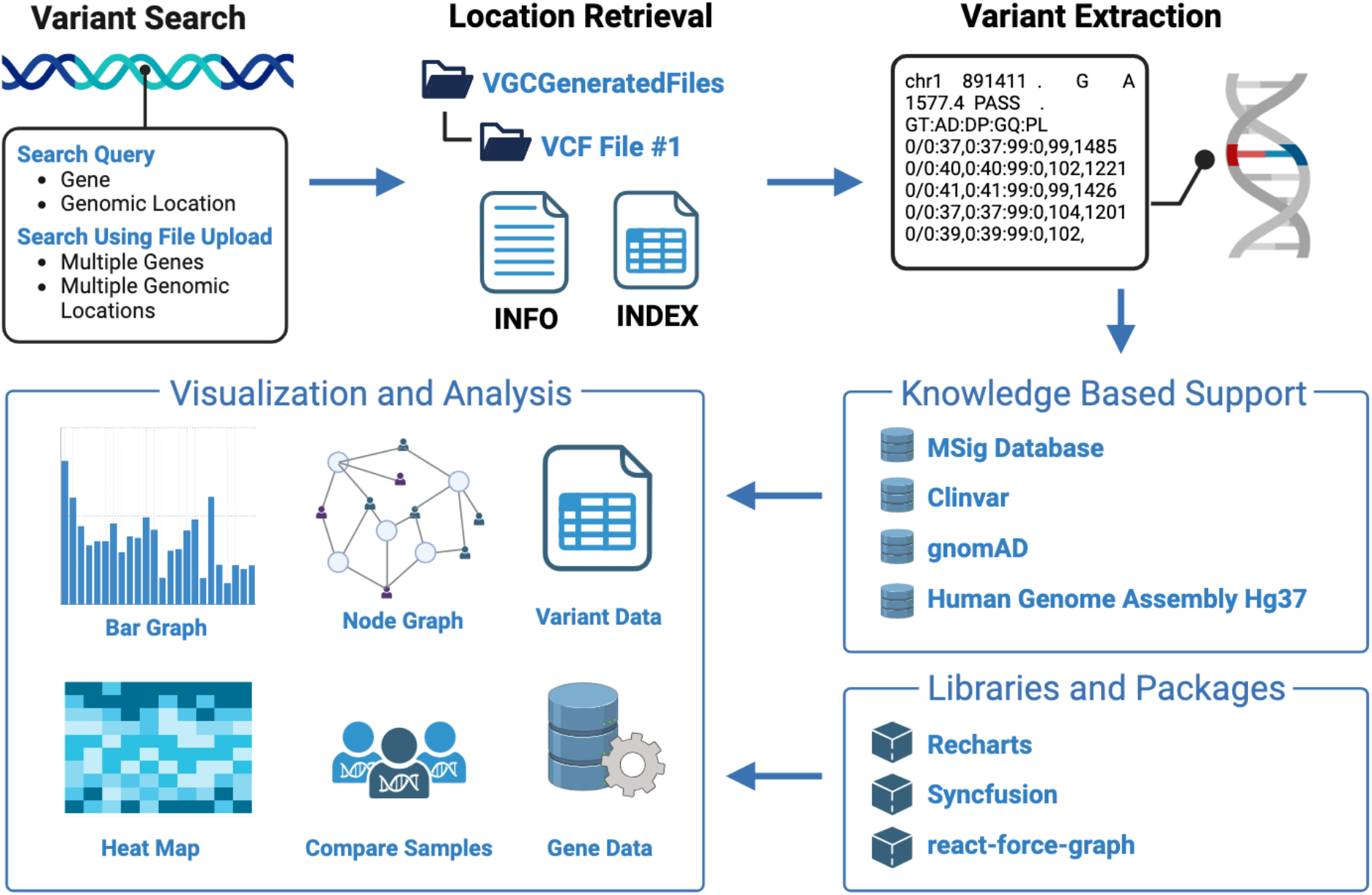
Design and integration of VGC. The query pipeline of VGC offers four distinct search options, as well as knowledge-based support with visualization and analysis.

### Programming languages, Applications and Libraries

VGC is a desktop application created using a JavaScript frontend and Java backend. The application is currently built using webpack module bundler version 5.86.0, and packed for iOS, Windows, and Linux using electron-forge.

Communication between the frontend and backend of VGC is handled by the Axios HTTP library. VGC is currently packaged using Electron for deployment, which allows the tool to be easily installed and run on a wide range of platforms and operating systems.

UI components are created using the React framework version 18.2.0, and styled using Tailwind CSS. To generate highly interactive and dynamic graphics for data visualization, the application utilizes a range of libraries, including Syncfusion, react-force-graph, and Recharts. These libraries provide a range of tools and functionalities for the visualization and analysis of complex data sets.

### Extraction and indexing of VCF

When a new VCF file is introduced to the program, VGC processes it to extract pertinent information, which is then stored in the user’s file system. A new directory named “VGCGeneratedFiles” is created in the user’s home directory, along with a corresponding directory that follows a specific naming scheme.

For each VCF file processed, a directory named “VGC_<filename>” is created. Inside these directories, two text files, named info_<filename> and index_<filename>, store important data. The info_<filename> file holds overall file information, such as the VCF file version, total number of samples, total number of chromosomes, number of variants, the header line, and a list of chromosomes in the file. The index_<filename> file contains chromosome-specific information. This indexing by VGC enhances response times for future queries. For each chromosome in the VCF file, the following details are listed: starting and ending lines, starting and ending positions, number of variants marked as “PASS,” and the count of pathogenic variants for that chromosome.

### Integration of Publicly Available Databases

VGC draws from a range of public databases, including MSig Database for GO terms, as well as KEGG, Biocarta, PID, and Reactome [9-13]. By leveraging these powerful databases, VGC is able to provide users with rich and detailed information about the genetic pathways and functions associated with their variant data, allowing for deeper insights and a greater understanding of the underlying biology. VGC also includes a dynamic link to gnomAD for variant information, allowing users to easily access and explore genetic variation data from this well-known database [14].

Additionally, the tool includes ClinVar data for pathogenic variant information, providing users with different visualization options for identifying and understanding potentially harmful genetic mutations [15]. VGC supports the Human Genome Assembly Hg37, ensuring compatibility with a wide range of data sets. The tool provides a range of options for exploring genetic variation, and can be tailored to the specific needs of the user by using optional phenotype input data.

### Dynamic Link to gnomAD for Variant Information

The dynamic link feature of VGC to gnomAD, a widely-used database for variant information provides users with a seamless connection to gnomAD, allowing them to access up-to-date and comprehensive variant data. By establishing this dynamic link, VGC ensures that users have access to the latest information on variant frequencies and population-specific data. This integration enhances the accuracy and reliability of variant interpretation, empowering researchers to make informed decisions based on the most current genomic data available.

### Incorporation of ClinVar Data for Pathogenic Variant Information

Inclusion of ClinVar data within VGC provides information on pathogenic variants and their clinical significance. By incorporating ClinVar data, VGC enables users to assess the potential pathogenicity of identified variants. Users can access curated information on variants that have been associated with specific diseases or conditions. This integration aids in variant prioritization, helping users focus on variants that may have clinical implications and guiding further investigation.

### Secure and Private Local Environment for Data Analysis

VGC is designed to run on the local machine or servers, ensuring that users can work with their genomic data in a secure and confidential setting. By avoiding the need to upload VCF files to the cloud, VGC protects sensitive genomic data and addresses privacy concerns. This local deployment approach instills a sense of reassurance in users, as they can confidently maintain control over their data, ensuring it stays within their organization’s infrastructure.

### Compatibility with Human Genome Assembly Hg37

VGC is designed to work seamlessly with this widely-used genome assembly, ensuring compatibility with a broad range of datasets. By supporting Hg37, VGC enables users to analyze genomic variation data generated using different platforms and datasets aligned to this assembly. This compatibility enhances the versatility and applicability of VGC, making it a valuable tool for a wide range of genomics studies and research projects.

### Customizing the Tool to Suit Individual User Requirements by Incorporating Optional Phenotype Input Data

VGC allows users to incorporate additional phenotype information, aligning the analysis with specific research questions or clinical contexts (Figure 2). By incorporating phenotype input data, VGC enables users to explore genetic variations in the context of specific phenotypic traits, enhancing the understanding of genotype-phenotype relationships. This customization feature makes VGC adaptable to various research and clinical scenarios, ensuring that users can leverage the tool to its full potential in their specific domain of interest.

**Figure 2:**
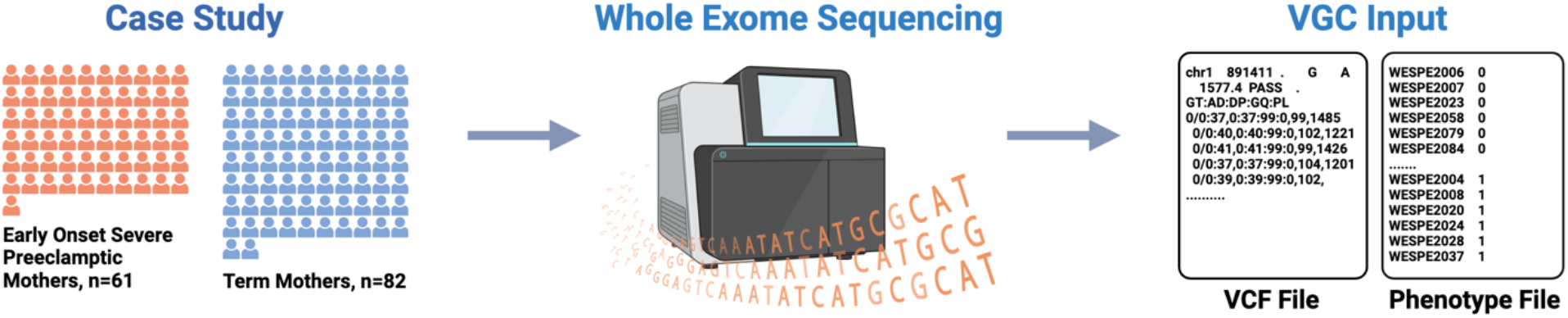
Schematic overview of case study to VGC input.

## Results

VGC features advanced visualization tools for VCF files. Demonstrating VGC’s capabilities, we present an example using whole exome sequencing data from preeclamptic patients and term mothers. The dataset includes 143 samples: 61 early onset severe preeclamptic cases and 82 term mother controls [16]. Through VGC, we offer a detailed analysis of this dataset, emphasizing major trends, statistical findings, and key outcomes aligned with our research goals. The insights gleaned from this study significantly enhance our understanding of variants associated with preeclampsia and offer valuable information for future research and practical applications.

### Comprehensive Variant Data Extraction and Visualization

VGC excels in variant browsing, offering features that enable effective exploration and analysis of genetic variations. It efficiently retrieves crucial data such as variant positions, alleles, genotype calls, and quality scores, offering a comprehensive and structured view of genomic variations for researchers and clinicians. For example, we demonstrate the visualization of variants in TTN, a gene with pathogenic, nominally significant variants identified in univariate analysis (refer to Figure 3). TTN variants are displayed in a histogram, sorted by variant position. Variants in intronic and exonic regions are differentiated by color. Users have the option to filter variants by categories such as “ALL,” “PASS,” or “Pathogenic”. VGC’s visualization capabilities extend beyond basic displays, offering sophisticated graphical representations that deepen understanding of variant data. Its intuitive and interactive visualizations allow users to discern patterns, connections, and insights within the genomic variations.

**Figure 3:**
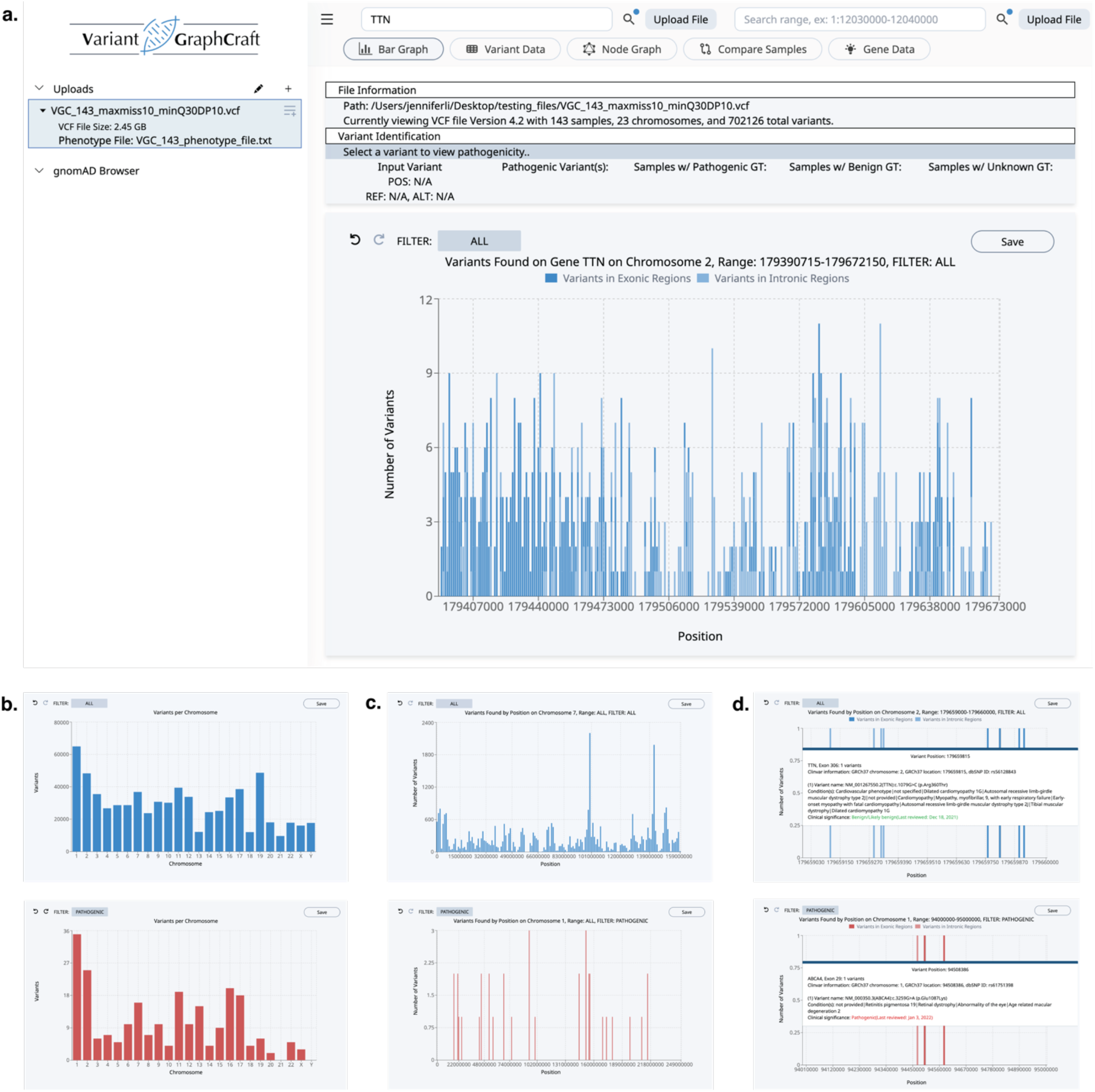
Histogram-based variant browsing with VGC. (a) The VGC user interface upon query of TTN, a gene found to contain pathogenic variants in the uploaded file. (b) Variants per chromosome, non-filtered [top] vs. filtered by pathogenicity [bottom]. (c) Partially magnified view of variants in CHR 1 for non-filtered [top] vs. filtered by pathogenicity [bottom]. (d) A detailed tooltip containing ClinVar-based information appears on hover when magnified to the single-position increment.

### Graph Representation of Samples and Genotype Data

VGC simplifies the interpretation of intricate genomic variation data by converting it into intuitive graphs, offering a visual summary of samples and their genotypes (as seen in Figure 4). By representing genotype data graphically, VGC enables users to effortlessly recognize patterns of genetic variation across different samples. This graphical format aids in exploring the relationships between genotypes, making it easier to identify common variants or unique genetic patterns within a population. Such a visual method enriches the users’ comprehension of the genetic landscape and assists in uncovering potential links between genotypes and phenotypes.

**Figure 4:**
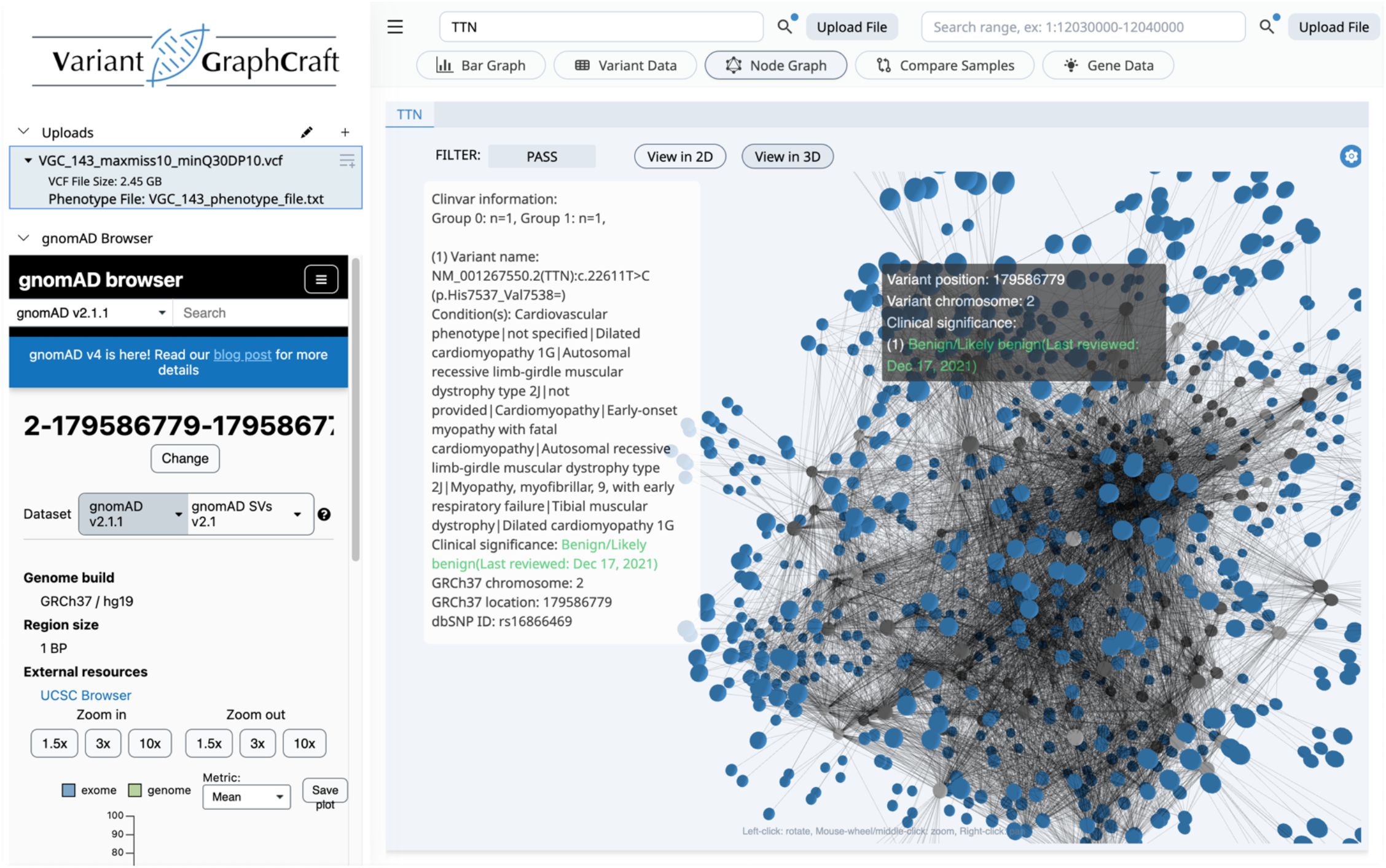
Force-graph visualization of variant to sample-grouping relations. Blue colored nodes show variants, while dark and light gray colored nodes represent cases and controls.

## Discussion

The features of VGC provide a comprehensive solution for users to easily analyze and visualize genomic variation data in a fast and secure manner. One key advantage of the tool is its user-friendly interface, which allows users to easily navigate and analyze large datasets. Another noteworthy feature is the fast filtering of millions of variants, which is crucial for researchers dealing with large-scale genomic data. This feature ensures that users can quickly identify the most relevant variants for further analysis. After initial upload of VCF files, even large files can be visualized in seconds in the next sessions. The ability to add and query based on any number of user-defined groups (or phenotypes) is a significant advantage for researchers interested in studying specific groups of individuals or genes. This feature allows for more targeted analysis. The tool’s ability to save and reuse analysis plans for reproducible research is a significant advantage, as it enables researchers to easily reproduce previous analyses and compare results. This feature is particularly important for ensuring that research findings are robust and reliable. The rapid VCF file browsing feature, with support for multiple visualizations such as histograms, spreadsheets, node graphs, and heatmaps, provides users with a comprehensive understanding of their data. This feature is particularly useful for identifying patterns and trends in genomic variation data. The tool’s ability to query by gene, range, position, and file upload, provides users with a range of options for searching and analyzing their data. This feature is particularly useful for identifying specific variants of interest and studying their potential impact on health and disease. The rapid identification and visualization of variant pathogenicity based on ClinVar data is another key advantage of VGC. This feature allows researchers to quickly identify potentially disease-causing variants, which can be further investigated for their clinical significance. VGC’s ability to display variant-to-sample genotype relations of user-defined groups is a significant advantage for researchers interested in studying the relationship between specific genetic variants and phenotypic traits. This feature allows for more targeted analysis and may lead to more insightful findings. The integrated variant querying through gnomAD, MSigDB, and Clinvar databases provides users with access to a wealth of public data, which can be used to enrich their own analysis. This feature is particularly useful for identifying novel variants and potential disease-causing mutations. Finally, the software’s design to run specifically on the local machine, with no VCF uploads to the cloud, ensures that users can work with their data in a secure and private environment. This feature is particularly important for researchers dealing with sensitive data and ensures that their research is conducted in a safe and confidential manner.

## Conclusion

In conclusion, the available features of VGC provide a comprehensive solution for researchers dealing with genomic variation data. The user-friendly interface, fast filtering, and ability to query based on user-defined groups, make it an efficient and effective tool for identifying potentially disease-causing variants. The ability to save and reuse analysis plans, rapid VCF file browsing, and integrated variant querying through public databases, further enhance the software’s capabilities, making it a valuable resource for genomic research. The tool’s rapid VCF file browsing with histogram, spreadsheet, node graph, and heatmap support further enhances its usability.

## Data Availability

VGC is freely available https://github.com/alperuzun/VGC

## Acknowledgements

We would like to thank Professor Vasileios P. Kemerlis of the Department of Computer Science at Brown University for his invaluable advice on security in developing Variant Graph Craft.

